# Synthetic Fibrous Hydrogels as Minimal Systems to Modulate Cell Migration Modes in 3D

**DOI:** 10.64898/2026.05.29.728729

**Authors:** Haoxiang Zhang, Guillermo Solís Fernández, Boris Louis, Sarah Vorsselmans, Johan Hofkens, Paul Kouwer, Hongbo Yuan, Susana Rocha

## Abstract

Cell migration in three-dimensional (3D) environments is highly plastic and regulated by extracellular matrix (ECM) cues. Engineered biomaterials provide controllable platforms to investigate how specific matrix signals regulate cell behavior in 3D, yet how defined biochemical signals control migration modes remain unclear. Here, we present tunable fibrous polyisocyanide (PIC) hydrogels functionalized with integrin-binding RGD peptides, cadherin-mimetic HAVDI peptides, or no ligands to direct mesenchymal, hybrid, or amoeboid-like migration of human adipose-derived stem cells without altering matrix mechanics. Using live-cell tracking, 3D displacement microscopy, matrix remodeling analysis, and YAP nuclear localization, we show that ligand identity governs adhesion organization, force transmission, and mechanotransduction. RGD-functionalized matrices promote β1-integrin clustering, extensive matrix remodeling, strong YAP activation and upregulation of migration-related genes. In contrast, non-adhesive matrices limit adhesion formation, resulting in weak force transmission and amoeboid-like behavior. HAVDI-functionalized matrices induce cadherin clustering and heterogeneous cellular responses, indicating that a hybrid migration mode arises from adhesion organization rather than a distinct transcriptional program. Together, these findings demonstrate that ligand identity alone is sufficient to program migration mode in a force-responsive 3D matrix and provide a versatile platform to dissect cell-matrix interactions in complex environments.

**Statement of significance:**

- Cell migration in tissues is highly adaptable, yet precise control of migration modes in defined 3D biomaterials remains challenging.
- We introduce fibrous PIC hydrogels presenting RGD, HAVDI, or no adhesive ligands to bias human stem cells toward mesenchymal-like, hybrid, or amoeboid-like migration states.
- By linking ligand identity to adhesion organization, matrix remodeling, YAP mechanotransduction, and gene expression, this work provides a minimal platform to dissect and engineer 3D cell-matrix interactions

## 1. Introduction

Cell migration is fundamental to embryonic morphogenesis, immune surveillance, tissue repair, and many pathological processes such as fibrosis and cancer invasion[1-5]. In living tissues, cells navigate through the heterogeneous three-dimensional (3D) extracellular matrix (ECM) while continuously adapting their morphology, force generation, and adhesion strategies[6, 7]. A hallmark of migration in 3D is phenotypic plasticity. Cells can adopt distinct migration modes, classically described as mesenchymal and amoeboid, while also occupying intermediate or hybrid states[8-10]. Furthermore, they can transition between these modes in response to environmental cues.

Mesenchymal migration is typically associated with elongated morphology, protrusion-driven movement, and strong interactions with the extracellular matrix through integrin mediated adhesions[11]. In contrast, amoeboid migration is characterized by a rounded morphology, reduced dependence on integrin adhesion and proteolysis, and higher cortical contractility, which allows cells to squeeze through pre-existing matrix pores[12]. Importantly, these migration modes are not restricted to specific cell types. Instead, they represent context dependent strategies used by many cells, including stem, immune, and cancer cells[5, 13-15]. For example, melanoma cells can undergo a mesenchymal-to-amoeboid transition that is accompanied by the acquisition of stem-like features[15], and the tumor microenvironment may promote this transition as an adaptive route to metastasis[16]. Similarly, macrophages migrating in collagen matrices can switch between mesenchymal and amoeboid modes depending on matrix architecture[17]. Understanding and ultimately controlling how cells adopt and transition between migration modes is therefore central to both fundamental biology and translational efforts to regulate tissue remodeling and improve therapeutic design.

Much of our mechanistic understanding of cell migration has been built on two-dimensional (2D) substrates and microchannel-based assays[5, 18, 19]. These platforms established foundational concepts linking adhesion dynamics, actin polymerization, actomyosin contractility, and mechanotransduction to cell motility[20-23]. However, these systems do not capture the fibrillar architecture, pore-scale confinement, and heterogeneous adhesive environments that define migration in 3D tissues.

In natural 3D ECMs, cells interact with discrete fibrils rather than a continuous planar surface, and their migration is shaped by steric confinement, local matrix architecture, and the balance between biochemical adhesion and mechanical force transmission[24]. Through these interactions, cells remodel their surroundings both biochemically, for example by locally degrading matrix components via proteases, and mechanically, by deforming or reorganizing matrix structures using cell-generated forces[25, 26]. However, because these structural, biochemical, and mechanical factors are highly coupled in natural ECMs, it remains difficult to independently dissect their respective contributions to cell migration.

To address this challenge, engineered 3D hydrogels have been used as controllable model systems that enable independent tuning of matrix parameters and more precise investigation of cell-matrix interactions. Both protease-mediated and force-mediated matrix remodeling have inspired important classes of engineered 3D migration models. On one hand, many engineered 3D hydrogels have been designed to support protease-dependent migration, for example by incorporating MMP-cleavable linkers into synthetic networks to decouple bulk mechanics from enzymatic remodeling[27-29]. On the other hand, increasing evidence shows that cells can also migrate through 3D matrices using physical, protease-independent strategies[30, 31]. Together, these findings highlight that migration in 3D is governed not only by matrix degradability, but also by the mechanical permissiveness of the surrounding network.

However, despite these advances, it remains difficult to determine how defined adhesion interfaces influence migration in 3D. Existing platforms have been highly valuable for studying matrix degradability and mechanical permissiveness, but few materials combine precise control over the biochemical adhesion cues with a force-responsive 3D matrix that can mechanically interact with cells. As a result, how specific adhesive interactions bias cells toward mesenchymal, amoeboid, or intermediate migration states remain poorly understood.

Here, we address this gap using polyisocyanide (PIC) hydrogels, a synthetic fibrous scaffold that mimics key structural features of the ECM and responds dynamically to cell-generated forces[32-35]. Building on our previous work showing that PIC hydrogels recapitulate force-driven matrix remodeling in a highly biomimetic manner[36], we functionalized PIC molecules with either an integrin-binding RGD peptide (PIC-R), an N-cadherin-mimetic HAVDI peptide (PIC-H)[37], or no adhesive ligand (PIC-N, **Fig. 1**a). Together, these conditions define a minimal 3D model that captures distinct interface scenarios encountered in tissues: integrin-mediated cell-matrix adhesion, cadherin-like cell-cell interactions, and a non-adhesive reference condition. Using this platform, we show that human adipose-derived stem cells adopt distinct migration phenotypes depending on the adhesion interface (**Fig. 1**b). PIC-R promotes polarized, protrusion-driven mesenchymal-like migration, PIC-N biases cells toward rounded morphologies with bleb-associated amoeboid-like behavior and limited persistence, and PIC-H yields heterogeneous intermediate phenotypes consistent with hybrid migration states. Using this minimal system, we further dissect how defined adhesion cues regulate migration mode selection in 3D and how these states are linked to matrix remodeling and downstream mechanotransduction pathways. Overall, this work establishes a reproducible synthetic platform to determine how biochemical adhesion signals shape the mechanics of 3D migration and provides a general strategy to engineer microenvironments that program migration behavior.

**Figure 1.**
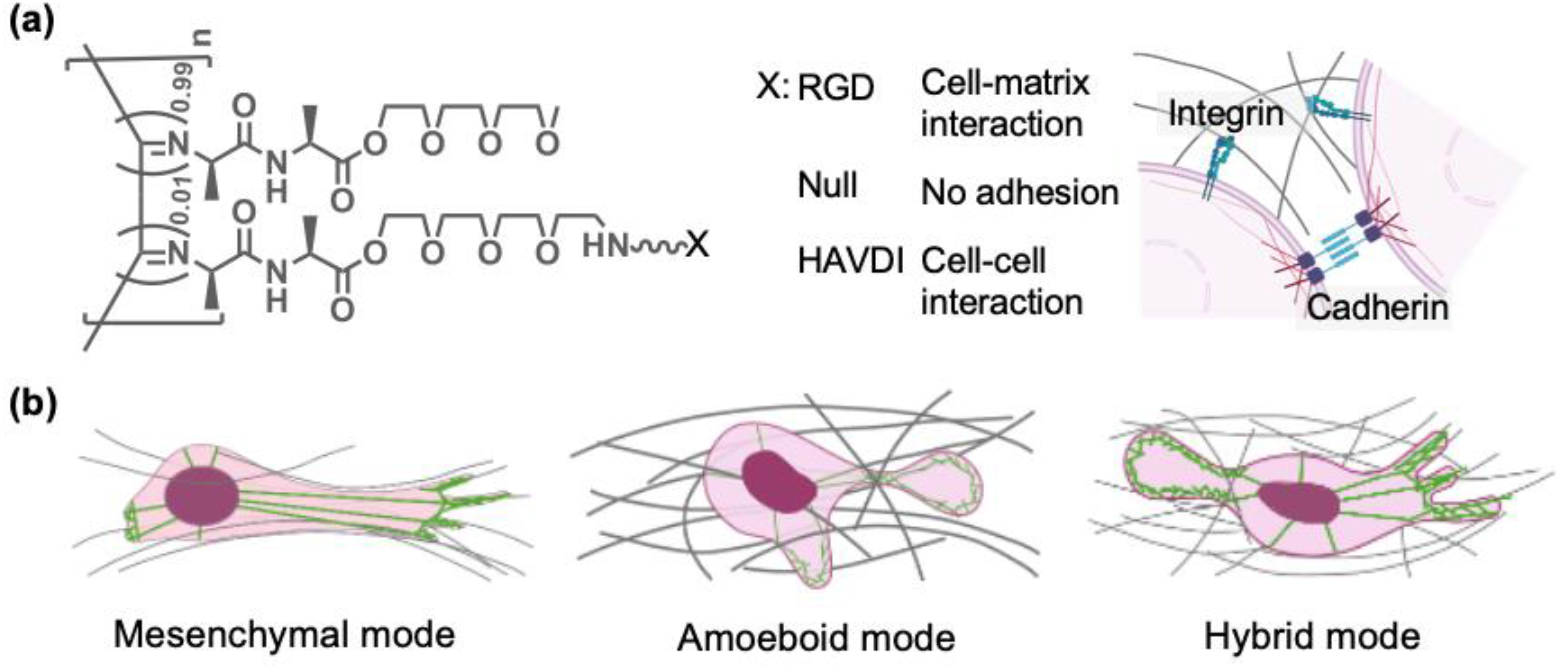
**(a)** Chemical structure of the PIC polymers used, showing the three different modifications (RGD, Null, HAVDI). RGD mimics integrin-mediated cell-matrix interactions, while HAVDI recapitulates cell-cell interactions. **(b)** Schematic illustration of the three different cell migration modes in 3D environments. The ECM fibers are depicted in grey, while the actin cytoskeleton is shown in green.

## 2. Material and methods

### 2.1 Polymer functionalization and hydrogel preparation

Polyisocyanides (PIC) were synthesized following an established protocol reported previously.[34, 36, 38] All polymers contained 3% azide-containing monomers. For biofunctionalization, linear RGD (promoting integrin binding) and HAVDI (N-cadherin-mimicking) peptides were used. The peptides were equipped with a DBCO-terminated spacer. The DBCO-PEG4-GRGDS (APi3844) and DBCO-PEG4-GGGGHAVDIG-OH (APi3845) peptides were obtained from AmbioPharm (Shanghai, China).

Peptide conjugation was performed via strain-promoted azide-alkyne cycloaddition following a previously described protocol[34]. Briefly, the azide-decorated polymer was dissolved in acetonitrile (2.5 mg mL^−1^), and the appropriate volume of DBCO-peptide solution in DMSO (12 mg mL^−1^) was added, resulting in approximately 1% ligand functionalization. The reaction mixture was stirred overnight at room temperature. Polymer-peptide conjugates were precipitated in isopropyl ether, collected by centrifugation, and air-dried for 2 to 3 days.

In the following text, RGD- and HAVDI-functionalized and non-functionalized PICs are referred to as PIC-R, PIC-H and PIC-N, respectively. Dry PIC polymers were sterilized by UV irradiation for 10 min and subsequently dissolved in PBS for 12 h at 4 °C to a final concentration of 2 mg mL^-1^. Polymer solutions were stored at 4 °C for no longer than one week prior to use in 3D cell culture experiments.

### 2.2 Cell culture and 3D encapsulation

Human adipose-derived stem cells (hASCs) were isolated from patients with informed consent, stored in the Radboud University Medical Center Urology Biobank (The Medical Ethics Review Committee of the Oost-Nederland Ethics Committee, approval CWOM9803-0060). The cells were cultured in Minimum Essential Medium Eagle (α-MEM) (Invitrogen, Thermo Fisher, USA). All media were supplemented with 10% fetal bovine serum (Sigma-Aldrich, USA) and 1% penicillin/streptomycin (final concentration of 100 IU mL^−1^ penicillin and 100 μg mL^−1^ streptomycin, Gibco, Thermo Fisher, USA).

Cells were harvested by trypsinization and resuspended in fresh culture medium. After cell counting, the cell suspension was mixed with the polymer solution on ice at a predefined ratio to obtain the desired cell density and polymer concentration. For cell migration experiments, the final cell density of 1 × 10^5^ cells mL^-1^ and a PIC concentration of 1 mg mL^-1^ were used, achieved by mixing equal volumes of a 2 mg mL^-1^ polymer solution and a 2 × 10^5^ cells mL^-1^ cell suspension. During mixing, pipetting was performed gently to minimize shear stress and prevent bubble formation.

The mixtures were transferred to containers and warm to 37°C to induce PIC gelation. After gel formation, prewarmed culture medium was added onto the samples. The samples were checked under microscope to confirm successful 3D cell encapsulation and were subsequently maintained in a standard cell culture incubator (humidified atmosphere, 37 °C, 5% CO2). Depending on the experiment, different containers were used, including glass-bottom Petri dishes (Cellvis, USA), 8-well chambered cover slides (Sigma-Aldrich, USA), or μ-Slide Angiogenesis plates (Ibidi, Germany).

### 2.3 Cell migration monitoring

Cells cultured in PIC hydrogels were monitored using a BioSPX Lionheart FX microscope equipped with 4× and 10× air objectives, under brightfield channels. The microscope incubator was set to 37°C and 5% CO2. Time-lapse images were acquired every 2 hours for a total of 60 hours. Image sequences were processed using ImageJ, where the image was aligned using the plugin ‘Linear stack alignment with SIFT’. Then individual cell trajectories were manually tracked with the ‘Manual Tracking’ plugin. The resulting coordinates were then replotted and analyzed using Python.

### 2.4 Fluorescence imaging of cellular components

For fluorescence imaging, μ-Slide Angiogenesis plates were used. Gels with encapsulated cells were washed three times with 1 mL prewarmed PBS and then fixed with 200 μL 4% paraformaldehyde (PFA) in PBS for 15 min in 37 °C. After fixation, the samples were washed with warm PBS, permeabilized with 0.1% Triton X-100 in PBS for 10 min at 37 °C, and subsequently blocked with 3% bovine serum albumin (BSA) in PBS for 1 h at 37 °C. For F-actin staining, samples were incubated with Phalloidin Atto-520 (1:1000 in 3% BSA/PBS, Sigma-Aldrich, USA) for 2-3 h. For immunostaining, samples were incubated with primary antibodies diluted in 3% BSA/PBS, including YAP antibody (63.7; sc-101199, Santa Cruz Biotechnology, USA), N-cadherin (13A9, sc-59987, Santa Cruz Biotechnology, USA), integrin β1(A4; SC-374429, Santa Cruz Biotechnology, USA) at 1:200 overnight at 37 °C. After washing with warm PBS, samples were incubated with the corresponding secondary antibody (Goat anti-Mouse IgG (H+L), Alexa Fluor 633; Invitrogen, USA) at 1:500 in PBS for 4 h at 37 °C. Samples were counterstained with DAPI (Thermo Fisher Scientific, D1306, 2.5 μg mL^−1^ in PBS) and washed with warm PBS.

All staining procedures were performed at a 37 °C heating plate and all the solutions used were pre-warmed to 37°C. This step is critical because PIC hydrogels are thermoresponsive, and insufficient temperature control can result in gel softening or liquefaction, causing sample loss during handling.

Fluorescence images were acquired on a Leica TCS SP8 X confocal microscope equipped with environmental control system to keep sample at 37°C, using water objectives (HC PL APO 25×/0.75, Leica and Fluotar VISIR 25×/0.95, Leica). For detection, we used a field-of-view scanner (400 Hz, bidirectional) and a hybrid photomultiplier detector (HYD-SMD, Leica). Transmission light detection was done in a separate channel using a standard Photomultiplier tube (Leica). Excitation was provided by a supercontinuum white light laser (470–670 nm, pulsed, 80 MHz; NKT Photonics) for Phalloidin Atto-520 and Alexa Fluor 633, and by a UV diode laser (405 nm, pulsed, 40 MHz; PicoQuant) for DAPI. Phalloidin Atto-520 was excited at 520 nm with emission collected between 530 and 700 nm. Alexa Fluor 633 was excited at 632 nm with emission collected between 639 and 773 nm, and DAPI was excited at 405 nm with emission collected between 410 and 551 nm.

Images were acquired around a range of 50 μm in the *z*-direction with a with a field of view of 77.6 × 77.6 μm^2^ (1024 × 1024 pixels). During measurements, A stage incubator was used to keep the cells were kept at 37 °C during image acquisition. The maximum projection images were prepared by compressing all the frames into one *z* projection using the LasX software (Leica Application Suite X V5.2.0).

### 2.5 3D matrix displacement microscopy

#### Image acquisition and processing

For displacement analysis, both cells and PIC hydrogel need to be stained. Adipose cells were incubated for 45 min with 12.5 µM CellTracker™ Green CMFDA (Life technologies, Belgium) in serum free medium before gel encapsulation. PIC polymers were labeled with DBCO-Atto647N to achieve fluorescent labelling of the fiber network. The volume ratio between DBCO-Atto647N: cell solution: polymer solution was 1:9:10. Briefly, the dye (10 μM) and the polymer solution (2 mg mL^-1^) were mixed on the ice first, and then the cold cell suspension (80,000 cells mL^-1^) was added and mixed gently. 12 µl of the mixture was pipetted in a µ-Slide Angiogenesis dish (Ibidi) and warmed up in an incubator.

#### Image acquisition

For PIC gel samples, fluorescence images were acquired using a Leica SP8 confocal microscope with a 25x water-immersion objective (NA 0.95) and a hybrid photomultiplier tube as detector (HYD-SMD, Leica). Cells labeled with CellTracker™ Green were excited at 492 nm, and PIC fibers grafted with Atto647N were excited at 646 nm with a non-sequential and bidirectional mode. The image stack was recorded with a distance of the confocal z-sections of 0.57 µm. With typically 177 z-slices and 512 x 512 pixels in the X-Y plane, the voxel dimensions were of 0.57 µm on the x, y and z direction. Images acquired under these conditions were defined as the *stressed state*, in which cells exert forces on the surrounding fibers, and were used for subsequent analysis.

Time-lapse image stacks were subsequently recorded after the addition of cytochalasin D (CytoD, 5 µM, Calbiochem®, Germany) at 20 min intervals between successive z-stacks. The stack acquired at the final time point was defined as the *relaxed state*, assuming that cellular contractile forces on the fibers were released following CytoD treatment. All other imaging parameters were identical to those used for the stressed state. A stage incubator was used to keep the cells were kept at 37 °C and 5% CO2 during image acquisition.

#### Displacement calculation

The Matlab toolbox TFMLAB[39] was used to process the microscopy data and calculate matrix displacements around the adipose cell[39, 40]. This software is available at: https://gitlab.kuleuven.be/MAtrix/Jorge/tfmlab_public. Briefly, the workflow of TFMLAB consists of three main steps: image processing, cell segmentation and displacement measurement.

##### Image processing

First, raw image data was filtered by penalized least squared-based denoising and enhanced with a contrast stretching operation. Second, stage drifts were corrected by applying rigid image registration with respect to the relaxed state. The shift was calculated by means of a phase correlation operation on the fibril images (i.e. acquired by means of fluorescence imaging for the PIC hydrogels and by means of second harmonic generation for the collagen gels). The calculated shift was then corrected for both the fibril images and the cell images.

##### Cell segmentation

Cell bodies were segmented by applying Otsu thresholding and by removing small binary objects

##### Displacement measurement and 3D visualization

TFMLAB uses the FFD-based image registration algorithm to register the stressed fibril images to the relaxed images. We used the normalized correlation coefficient as the similarity metric and a stochastic gradient descent method with adaptive estimation of the step size as the optimizer[41]. As a result, a 3D displacement field vector was obtained at every voxel of the image. The software Paraview 5.13.3 was used to create 3D renders of the cells and the displacement field using the .vtk files provided by TFMLAB.

### 2.6 Matrix remodeling analysis

#### Image acquisition and processing

Fluorescence images were acquired using the same samples and imaging settings as those used for the stressed state in the matrix remodeling experiments. Cells were labeled with CellTracker™ Green, and PIC fibers were grafted with Atto647N. Images were subsequently processed using a custom MATLAB code (*FiberRemodeling*), as described below.

#### Cell segmentation

The cell was segmented by applying Otsu thresholding and by removing small binary objects.

#### Polymer segmentation

The polymer was segmented by applying an intensity threshold based on the 99th percentile intensity of the pixels on 10% outside frame of the image. Small objects were then removed, and small holes filled.

#### Calculations

The volume was calculated by summing all pixels inside the polymer segmented mask multiplied by the voxel size and subtracting the cell volume from it. The distance between the cell and polymer was calculated as follows: for each point on the cell mask surface, the closest point on the polymer mask surface (outer shell) was determined using Euclidean distances. We then looked at the distribution of those distances to have an estimate on how far the densified region can extend from the cell.

#### 3D Modeling

The 3D models of cell and polymer were made by plotting the surface for i=0.5 which created a surface at the surface of the segmented mask.

#### Code source

All the code related to the matrix remodeling analysis can be found at: https://github.com/BorisLouis/fiberRemodeling

### 2.7 3D YAP nucleus-to-cytoplasm ratio analysis

The 3D nuclear-to-cytoplasmic (N/C) ratio of YAP was quantified using a custom analysis pipeline developed in Python. Z-stack fluorescence images containing DAPI (nuclear) and YAP channels were imported directly from Leica LIF files, preserving the original image resolution and intensity information.

Nuclear regions were segmented from the DAPI channel using automated thresholding on denoised images, followed by morphological filtering to remove small artifacts and to retain biologically relevant nuclear structures. The cellular region was defined from the YAP channel using automated intensity-based thresholding, with subsequent morphological refinement to delineate the full cell area. The cytoplasmic region was defined as the cellular mask excluding the nuclear mask. In cases where multiple cells were present within a single imaging field, all segmented cell regions were included in the analysis.

For each z-slice containing a valid nuclear region, the YAP N/C ratio was calculated as the mean fluorescence intensity of YAP within the nuclear region divided by the mean fluorescence intensity within the cytoplasmic region. To obtain a representative 3D YAP N/C ratio for each imaging field, a weighted average of the per-slice N/C ratios was computed, with weights proportional to the segmented cell area in each z-slice. A quality-control module was used to visualize segmentation results and quantitative metrics, enabling manual verification of nuclear and cytoplasmic regions and minimizing potential errors arising from automated image analysis. Fig. S3 All code related to the 3D YAP nuclear-to-cytoplasmic ratio analysis will be available by request.

The resulting weighted N/C ratio was taken as the final 3D YAP nuclear-to-cytoplasmic ratio for each sample. At least 25 independent imaging fields were analyzed per hydrogel condition.

### 2.8 Quantitative real-time PCR (qPCR)

#### RNA extraction and cDNA synthesis

For qPCR analysis, total RNA was isolated from cell pellets at 48 h after seeding hASCs. NZY total RNA isolation kit (NZYtech, Portugal) was used for total RNA isolation following the manufacturers protocol. After isolation, total RNA was further concentrated using the RNA Clean & Concentrator-5 kit (Zymo Research, USA) and then was quantified with a BioDrop spectrophotometer (Biochrom, UK). Then, cDNA was obtained from 400 ng of total RNA using NZY M-MuLV First-Strand cDNA Synthesis Kit (NZYtech, Portugal) according to manufacturer’s instructions. Separate oligos cDNA synthesis kit with the oligodT oligos following the manufacturer’s protocol. ***qPCR reaction***. For qPCR analysis cDNA was diluted 1:10 and 2µL of the diluted cDNA were used for each qPCR reaction. Quantitative PCR was performed using Xpert Fast SYBR BLUE (Syntezza Bioscience, Israel), and PCR and data collection were performed on a Quant Studio 5 (ThermoFisher). All the oligos used can be found on **Table S1**.

#### Normalization and analysis

All quantifications were normalized following the ΔΔCt method, using human RPL19 as housekeeping control. The expression for each of the conditions was evaluated relative to the undecorated PIC-N encapsulated cells.

### 2.9 Data analysis

Statistical analysis and data plotting was performed with GraphPad Prism 10. Normality was assessed using the Shapiro–Wilk test. Differences between two groups were analyzed using unpaired two-tailed Student’s t-test or Mann–Whitney U test as appropriate. For comparisons among multiple groups, one-way ANOVA or Welch’s one-way ANOVA followed by Dunnett’s T3 multiple comparisons. Statistical significance is denoted by \**p* < 0.05, \*\**p* < 0.01, \*\*\**p* < 0.001, and \*\*\*\**p* < 0.0001.

## 3. Results and Discussion

### 3.1 Ligand identity shape migration phenotypes

To determine whether defined adhesive interfaces bias early cell behavior in 3D, we first examined cell morphology and cytoskeletal organization in PIC matrices presenting distinct ligands: PIC-R, PIC-H, and PIC-N. Human adipose derived stem cells (hASCs) were used as a model to examine how ligand identity influences cell-matrix interactions and related cell behaviors in 3D. Brightfield imaging 48 h after cell embedding revealed clear differences in cell morphology across the three conditions, which became more pronounced over time (**Fig. S1)**.

To resolve the underlying cytoskeletal organization, cells were fixed at 48 h, stained with phalloidin to visualize filamentous actin, and imaged by confocal imaging (**Fig. 2**a-f). In PIC-R, hASCs displayed pronounced spreading, with elongated cell bodies and well-defined protrusions (**Fig. 2**a,d). Actin formed prominent bundles aligned with the long axis of the cell and extended into protrusions, consistent with a polarized mesenchymal-like morphology. In contrast, cells in PIC-N remained largely rounded and showed predominantly cortical actin, with no robust actin bundles or stress fiber-like. Instead, only fine, diffuse actin surrounded the cell body, consistent with weak matrix coupling and low-adhesion in amoeboid-like behavior structures (**Fig. 2**b,e). Interestingly, PIC-H yielded the broadest distribution of morphologies. Although most cells exhibited limited spreading, they frequently formed short protrusions and occasional membrane bleb-like structures (**Fig. 2**c,f). Actin in these protrusions was shorter and less aligned than in PIC-R, suggesting reduced actin bundling and a distinct mode of force organization, indicating of an intermediate cellular state between amoeboid and mesenchymal phenotypes.

**Figure 2.**
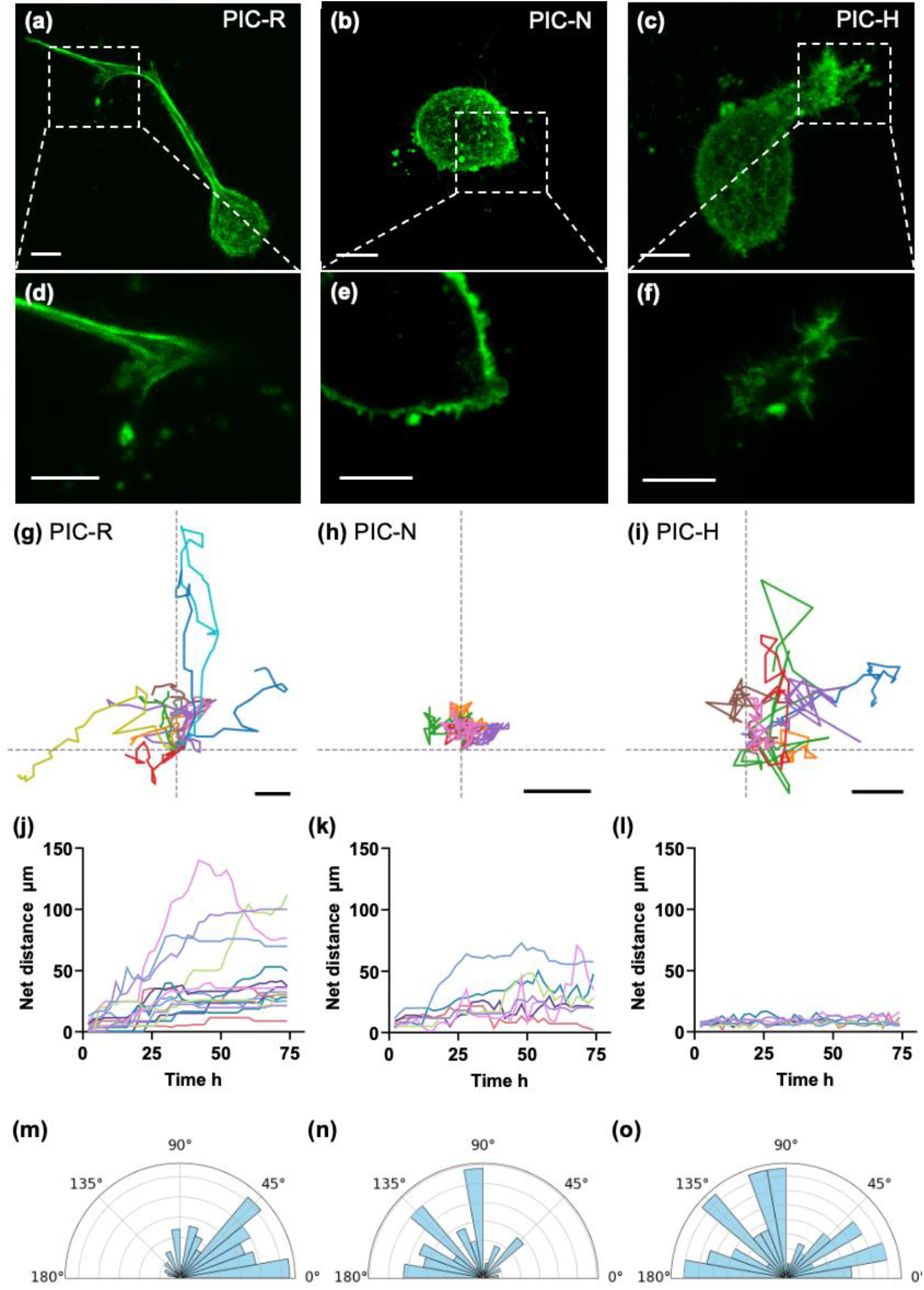
Ligand-dependent cell morphology and migration in 3D PIC hydrogels. **(a-f)** Representative fluorescence images of adipose derived stem cells encapsulated in PIC-R **(a, d)**, PIC-N **(b, e)** and PIC-H **(c, f).** Filamentous actin was stained using fluorescently-labelled phalloidin. **(a-c)** Maximum projection of the z-stacks images; scalebar: 25 μm. **(d-f)** Magnification of the area indicated by the white square in **(a-c)**; only a single xy-image of the z-stack is shown; scalebar: 10 μm; **(g-i)** Representative trajectories described by single cells in PIC-R **(g)**, PIC-N **(h)**, and PIC-H **(i)**. The trajectories correspond to the movement of the cells shown in Figure S1 and were aligned to a common origin at the starting position for easier comparison; scalebar: 20 µm. **(j-l)** Net displacement of the different individual trajectories, from the starting position, as a function of time, for cells migrating in PIC-R **(j)**, PIC-N **(k)** and PIC-H **(l). (m-o)** Rose plots showing the distribution of turning angles between consecutive time points of each individual trajectory, for cells migrating in PIC-R **(m)**, PIC-N **(n)** and PIC-H **(o)**.

Building on the distinct morphological phenotypes described above, we next investigated how matrix functionalization influenced cell migration behavior in 3D. Time-lapse brightfield imaging of hASCs was performed over a time course of 60 hours and the subsequent migration trajectories obtained for the cells were analyzed (**Fig. 2**g-o). Cells embedded in PIC-R exhibited persistent, directional migration, with sustained translocation over longer distances and showed a clear front-rear polarity while migrating (**Fig. 2**g). This behavior is in agreement with the spindle-like morphology and aligned actin organization observed in PIC-R, which are typical features of mesenchymal like migration in 3D matrices. In contrast, cells in PIC-N showed shorter, spatially confined trajectories and frequently reversed travelling direction, resulting in limited net translocation (**Fig. 2**h). These features are consistent with the rounded morphology and dynamic, bleb-like protrusions observed in this condition, indicative of low-adhesion amoeboid-like migration. Cells in PIC-H displayed intermediate migration behavior, with moderate displacement, frequent changes in direction and continuous shape fluctuations during movement (**Fig. 2**i). Migration was not consistently coupled to a single protrusion axis, in line with the heterogeneous morphology and mixed protrusive features observed, suggesting a mixed or intermediate migration mode. Comparable migration phenotypes have previously been referred to as amoeboid-filopodial[42], hybrid mode[10] or hybrid amoeboid-mesenchymal[8].

Quantitative trajectory analysis supported these observations (**Fig. 2**j-o). Net displacement from the starting position increased fast in PIC-R, reached intermediate values in PIC-H, and remained low in PIC-N (**Fig. 2**j-l). Turning behavior showed the opposite trend: cells in PIC-N exhibited the largest turning angles, while PIC-H showed intermediate values, and PIC-R displayed the smallest angles, indicating the highest directional persistence in PIC-R (**Fig. 2**m-o). Together, these results demonstrate that, within a synthetic fibrous scaffold of comparable bulk mechanics, the identity of the adhesion interface is a key determinant of migration persistence and effective translocation in 3D confined materials.

### 3.2 Adhesion cues recruit distinct molecular systems

Cell migration requires coordinated organization of the cytoskeleton and adhesion-related components, which together support protrusion formation, force transmission, and directional movement. Based on the distinct cell morphologies and migration behaviors observed across the three PIC matrices, we hypothesized that the different ligands presented by the materials. promote the formation of distinct adhesion structures. To investigate ligand-specific adhesion organization at the subcellular level, we examined the organization of key adhesion proteins by immunostaining. Since PIC-R and PIC-H present ligands that engage integrin- and cadherin-based adhesions, respectively, we examined the subcellular organization of integrin β1 and N-cadherin (**Fig. 3**). In PIC-R, integrin β1 staining revealed prominent punctate signals at the cell membrane, especially near the tips of cellular protrusions (**Fig. 3**a,d), consistent with integrin clustering at adhesion sites in cells engaging with the RGD-functionalized matrix. In contrast, PIC-H and PIC-N showed more diffuse integrin β1 staining, with no comparable punctate organization, suggesting limited integrin clustering in these conditions (**Fig. 3b,c,e,f**). When examining N-cadherin staining, PIC-H samples displayed prominent punctate membrane-associated clusters (**Fig. 3**i,m), consistent with localized HAVDI-mediated adhesion. N-cadherin signal was detected in PIC-R or PIC-N conditions (**Fig. 3**g,h,j,l).

**Figure 3.**
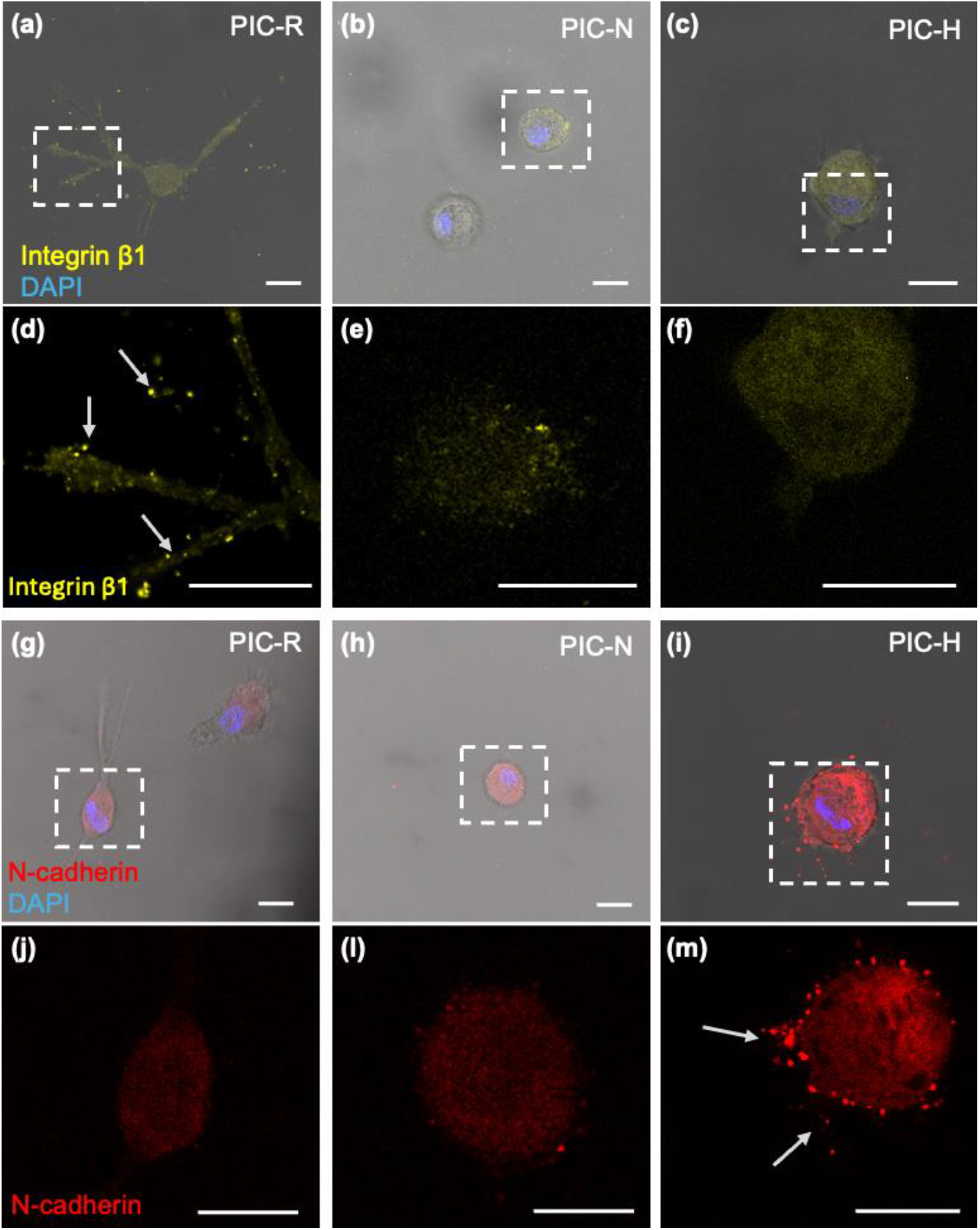
Ligand-specific clustering of integrin β1 and N-cadherin in PIC hydrogels. **(a-c)** Representative overlay images of brightfield and integrin staining (yellow: integrin, blue: DAPI, gray: brightfield), of adipose derived stem cells embedded in PIC-R **(a)**, PIC-N **(b)** and PIC-H **(c)**; **(d-f)**, Magnification of the area indicated by the white square in **(a-c)**, showing only the integrin signal. **(g-i)** Representative overlay images of brightfield and cadherin staining (red: N-Cadherin, blue: DAPI, gray: brightfield), of cells embedded in PIC-R **(g)**, PIC-N **(h)** and PIC-H **(i)**; **(j-m)** Magnification of the area indicated by the white square in **(g-i)**, showing only the cadherin staining. Scale bar: 25 μm.

Together, these results demonstrate that PIC functionalization promotes ligand-specific subcellular organization of adhesions. RGD functionalization (PIC-R) supports integrin clustering and is associated with robust actin bundling and polarized morphology, consistent with mesenchymal-like migration. HAVDI functionalization (PIC-H) selectively enriches N-cadherin at the cell periphery and coincides with heterogeneous cell shapes and intermediate migration behavior. In PIC-N, the absence of adhesive ligands is accompanied by weak recruitment of adhesion markers and a rounded morphology. These ligand-dependent adhesion patterns provide a mechanistic basis for the distinct migration behavior and matrix interaction phenotypes observed across the three conditions.

### 3.3 Ligand identity tunes matrix coupling and remodeling

Because adhesive ligands function as the interface that orchestrates cell-hydrogel interactions, we hypothesized that ligand-specific adhesion complexes in PIC-R and PIC-H would enable cells to mechanically couple to the surrounding network and transmit forces into the matrix. To test whether distinct cell-hydrogel interactions lead to different levels of mechanical coupling, we quantified cell-induced matrix displacements in the three materials using 3D matrix displacement microscopy. Briefly, following a previously established approach[36], PIC polymers were fluorescently labeled to visualize their fiber network structure (**Fig. 4**a,b). Embedded cells were relaxed by pharmacological disruption of actomyosin contractility using cytochalasin D (CytoD, **Fig. 4**c). Changes in fiber position before and after relaxation were subsequently used to reconstruct the three-dimensional displacement field around individual cells from the difference between these two states (**Fig. 4**d-f).

**Figure 4.**
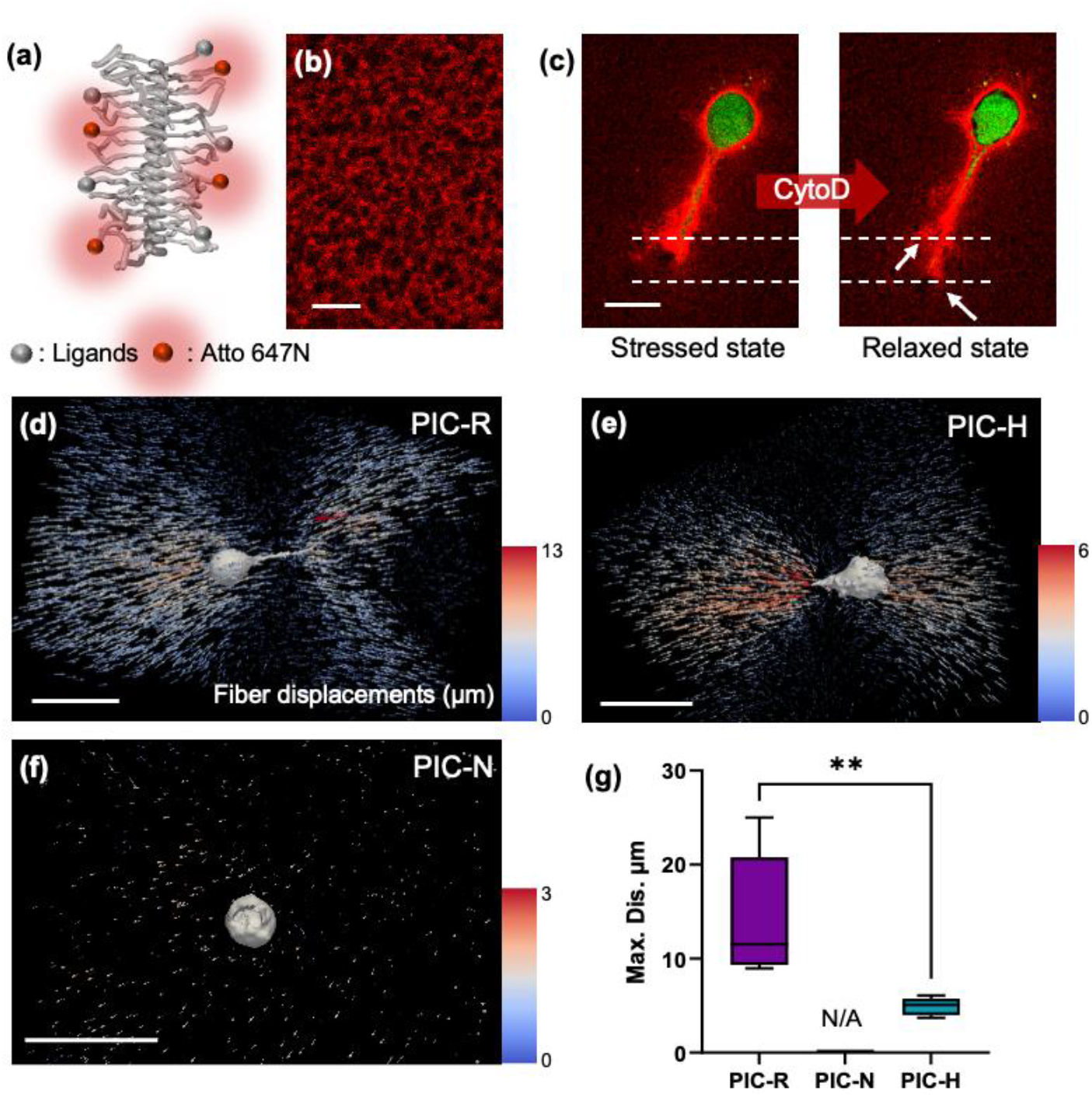
Ligand-dependent fiber displacements in PIC hydrogels. **(a)** Schematic representation of Atto647-labeled PIC polymers. **(b)** Representative fluorescence image of PIC-N, displaying the fibrous network; scalebar: 10 μm; **(c)** Representative images used for displacement analysis showing the PIC network in the stressed state and relaxed state (after addition of CytoD). The cell is fluorescently labelled using CellTracker. White arrows indicate regions of large fiber displacement. Scalebar: 25µm **(d-f)** Representative displacement fields around single cells in PIC-R (d), PIC-H (e), and PIC-N (f). Vector color indicates displacement magnitude in micrometers. Scalebar: 50µm (g) Maximum displacement magnitude per cell quantified across conditions (N = 5). Differences between two groups were analyzed using unpaired two-tailed Student’s t-test.

In PIC-R, cells generated pronounced displacement fields that extended from the cell body into the surrounding network (**Fig. 4**d). Displacement vectors were spatially continuous and reached high magnitudes, consistent with strong mechanical coupling between the actin cytoskeleton and the fibrous matrix through integrin mediated adhesion. In PIC-H, displacement fields were also detectable, but their magnitudes were consistently lower, suggesting that HAVDI-mediated N-cadherin-hydrogel interactions support force transmission into the matrix, albeit less efficiently than RGD-mediated coupling (**Fig. 4**d). In PIC-N, displacement fields were minimal and close to background levels, consistent with the limited formation of stable adhesion complexes (**Fig. 4**e). Together, these results show that the identity of the adhesion interface strongly regulates the magnitude and spatial extent of cell-induced matrix displacements, providing a direct mechanical correlation with the different cellular morphologies and migration phenotypes observed.

Beyond force transmission, cells migrating in 3D environments physically remodel the surrounding matrix. Such remodeling can alter the local microenvironment and is expected to depend on both the identity of the adhesion ligand and on the efficiency of the mechanical coupling between the cell and the fibrous network. To quantify these effects, we imaged fluorescently labeled PIC fibers together with labeled cells by confocal microscopy and analyzed pericellular remodeling using a three-dimensional segmentation workflow (see Methods, section 2.6). This approach enabled extraction of both cell volume and adjacent regions of higher fiber signal intensity that reflect local matrix reorganization. As shown in **Fig. 5**, cells in PIC-R and PIC-H were surrounded by regions of increased PIC fluorescence intensity, indicating local fiber densification around the cell body and along protrusions. In PIC-R, these densified regions were larger and more continuous, frequently extending along the full length of protrusions (**Fig. 5**d). In PIC-H, remodeling was detectable but more localized and variable between cells, typically appearing as discrete patches near the cell surface (**Fig. 5**f). In contrast, cells in PIC-N showed minimal evidence of fiber densification. Instead, regions of reduced fluorescence signal were frequently observed adjacent to cells. (**Fig. 5**e). Since PIC is non degradable for cells, this likely reflecting local displacement or reorganization driven by amoeboid-like motion rather than adhesion-mediated fiber remodeling.

**Figure 5.**
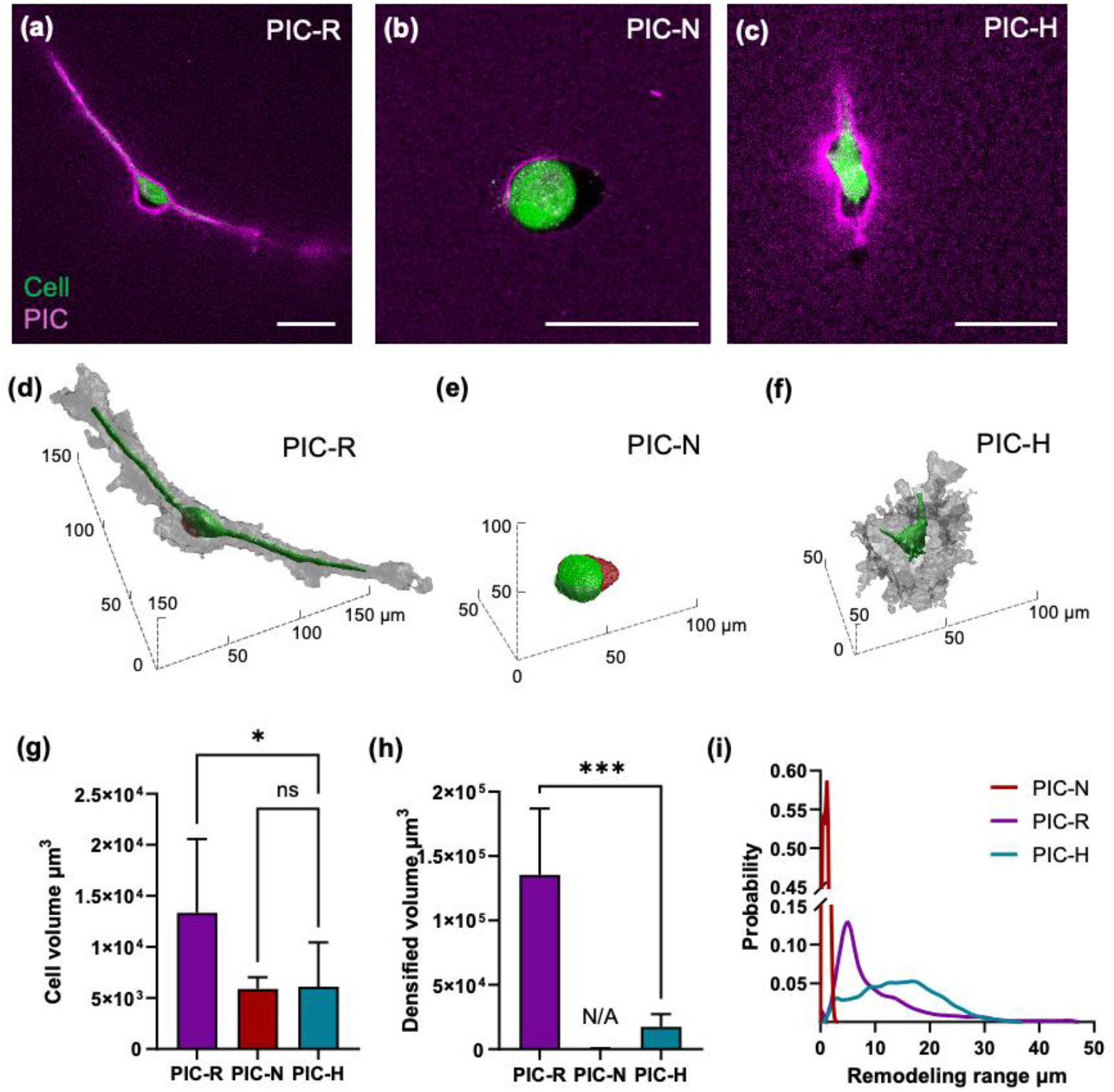
Cell induced fiber remodeling in the different PIC hydrogels. **(a-c)** Representative fluorescence images of cells in embedded in PIC-R (a), PIC-N (b) and PIC-H(c); magenta: PIC network, green: cell, Scalebar: 50 μm; **(d-f)** 3D rendering of the densified volume arising from fiber remodeling; gray: densified volume, green: cell, red: dark volume around the cell. **(g)** Cell volume of cells embedded in different hydrogels. Differences between two groups were analyzed using one-way ANOVA. **(h)** Volume of densified region surrounding the cells. Differences between two groups were analyzed using Mann– Whitney U test. **(i)** Probability density distribution of remodeling range. (N > 5)

Three-dimensional rendering confirmed these trends and enabled quantitative comparison (**Fig. 5**d-i). Cells exhibited the largest volume in PIC-R, with smaller measured volumes in PIC-H and PIC-N (**Fig. 5**g). Consistent with the qualitative observations, the volume of densified fibers was substantially higher in PIC-R than in PIC-H, while PIC-N showed negligible densified volume (**Fig. 5**h). Analysis of remodeling range distributions revealed that RGD-mediated adhesion supports remodeling over larger spatial distances than HAVDI-mediated adhesion, in line with stronger and more extensive force transmission into the fibrous network (**Fig. 5**i).

Together, these results demonstrate that ligand identity strongly regulates the magnitude and spatial organization of pericellular matrix remodeling. RGD-mediated integrin engagement promotes extensive and continuous densification, whereas HAVDI-mediated interactions support weaker and more heterogeneous remodeling. The absence of adhesive ligand largely abolishes densification. These differences reflect distinct mechanical strategies employed by cells under different adhesive constraints. Specifically, the extensive and continuous densification observed in PIC-R is consistent with mesenchymal migration where cells generate tension in the surrounding matrix through contractility to facilitate forward movement [43]. This structural reorganization of the fibrous network likely enhances force transmission and supports persistent migration in 3D.[25, 44, 45]

### 3.4 Distinct ligands drive differential intracellular states

To further investigate whether the distinct adhesion complexes and migration behaviors observed across the three PIC matrices are associated with differences in intracellular mechanosensing, we analyzed the subcellular localization of YAP, a mechanosensitive transcriptional coactivator[46-48]. hASCs were co-stained for YAP and DAPI, and the 3D nuclear-to-cytoplasmic ratio of YAP intensity was quantified from fluorescence images at 4, 12, and 48 h after cellular embedding. For easier comparison between the fluorescence intensity in the nucleus and cytoplasm, nuclei contours calculated from the DAPI images are shown as white lines in **Fig. 6**.

**Figure 6.**
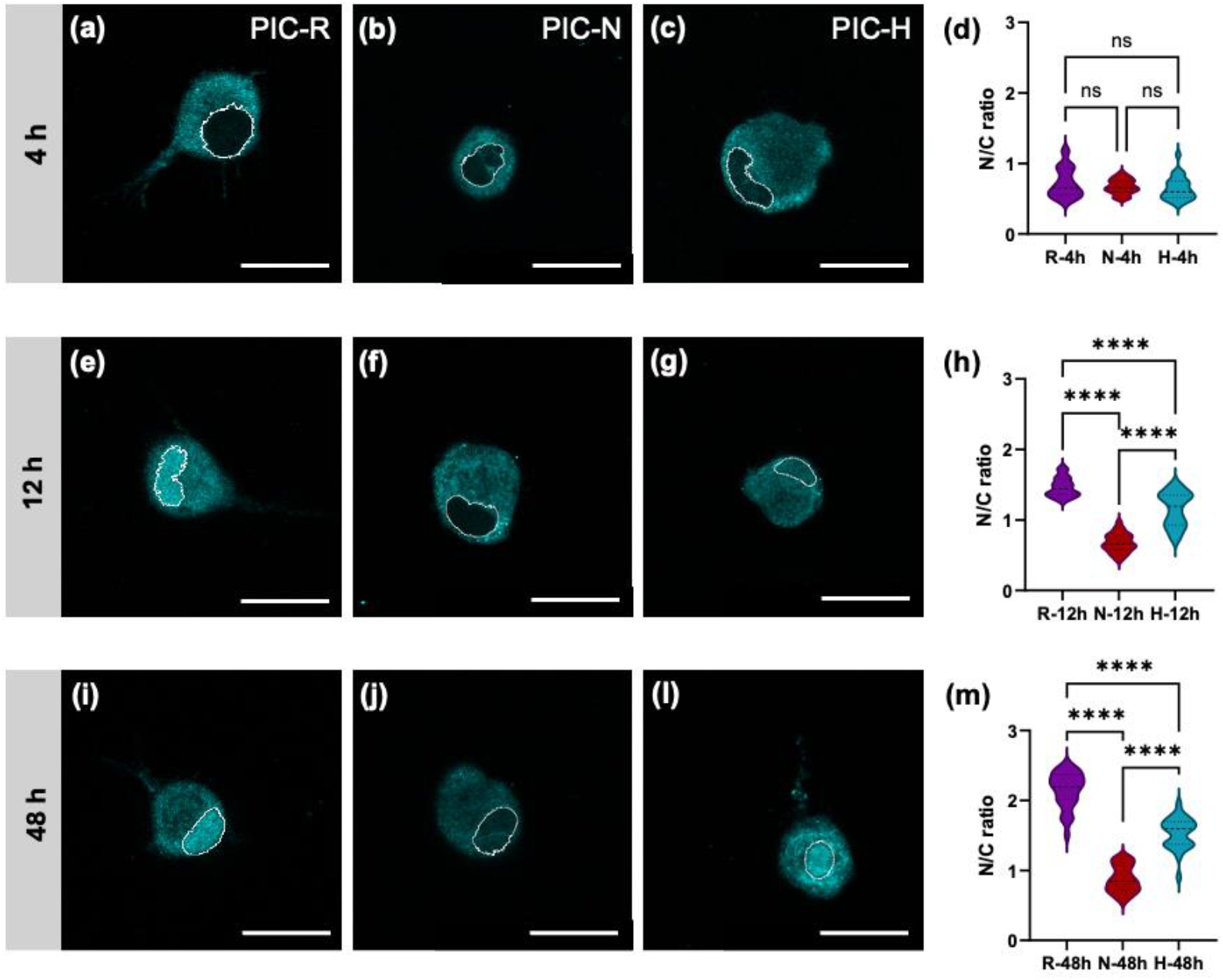
Ligand-dependent YAP nuclear localization in PIC hydrogels. **(a-c, e-g, i-l)** Representative fluorescence images of YAP in cells embedded in PIC-R (a, e, i), PIC-N (b, f, j) and PIC-H (c, g, l), at 4 (a-c), 12 (e-g) and 48h (i-l). The white like indicate the contour of the nucleus, determined using a nuclear stain (DAPI); details in the Methods section. Scalebar: 25 μm **(d, h, m)** Nucleus-to-cytoplasm (N/C) ratio of YAP intensity, calculated from confocal z-stacks, for the different time points. (N > 26). Statistical significance was determined using Welch’s one-way ANOVA with Dunnett’s T3 multiple comparisons test.

At 4 h, YAP nuclear-to-cytoplasmic ratios were comparable across PIC-R, PIC-N, and PIC-H, indicating that early after embedding, cells had not yet established distinct mechanotransduction states (**Fig. 6**a-d). By 12 h, clear differences emerged between conditions (**Fig. 6**e-h). Cells in PIC-R showed significantly higher YAP nuclear localization than cells in PIC-N, consistent with stronger force transmission associated with integrin-mediated matrix engagement. PIC-H exhibited intermediate YAP localization, with broader distributions, indicating increased cell-to-cell variability (**Fig. 6**h). At 48 h, these trends persisted, with PIC-R maintaining the highest YAP nuclear enrichment, PIC-N remaining lowest, and PIC-H displaying intermediate and heterogeneous responses (**Fig. 6**i-m).

YAP nuclear translocation is widely used as a readout of mechanical signal transmission. These findings indicate that ligand identity modulates mechanotransduction through distinct modes of mechanical coupling. In PIC-R, integrin-mediated adhesions are likely coupled to actin bundles and force-generating structures that efficiently transmit tension to the nucleus, promoting robust YAP nuclear localization. In PIC-H, cadherin-mediated interactions may preferentially organize peripheral and cortical actin structures that transmit lower and more variable tension, leading to intermediate and heterogeneous YAP responses. In PIC-N, limited formation of adhesive complexes restricts stable force transmission, and confinement alone appears insufficient to drive YAP accumulation in the nucleus. Together, these findings link ligand-dependent migration and matrix remodeling phenotypes to differences in intracellular force transmission and nuclear mechanotransduction.

### 3.5 Ligand-dependent adhesion modulates gene expression

Since YAP is a mechanosensitive transcriptional coactivator, the ligand-dependent differences in force transmission and YAP localization observed in the previous sections may ultimately influence gene expression programs linked to cell migration and matrix interaction. To further investigate whether the distinct migration behaviors observed in the three PIC matrices are associated with differences in downstream transcriptional regulation, we analyzed the expression of migration-related genes.

As a starting point, we leveraged a previously published proteomic dataset comparing cells cultured in PIC matrices functionalized with RGD or lacking adhesive ligands[49] . Here, we selected a subset of migration- and adhesion-related candidates from this dataset and quantified their expression by qPCR in the current system, using the same cell type (hASCs) but under the present experimental conditions (**Table 1**).

**Table 1.** Genes of migration related proteins.

| Gene | Full name | Over express in<sup>1</sup> | Function category |
| --- | --- | --- | --- |
| **CYR61** | Cysteine-rich angiogenic inducer 61 | RGD | ECM & adhesion |
| **TSP1** | Thrombospondin-1 | RGD | Matrix remodeling |
| **FBLN2** | Fibulin-2 | RGD | ECM organization |
| **MXRA5** | Matrix remodeling-associated protein 5 | RGD | ECM remodeling |
| **GPNMB** | Transmembrane glycoprotein NMB | N3 | Adhesion / signaling |
| **GAP43** | Neuromodulin | N3 | Cytoskeleton dynamics |
| **CHI3L1** | Chitinase-3-like protein 1 | N3 | Stress/ECM regulation |
| **ITGA2** | Integrin $\alpha 2$ | N3 | Cell-ECM adhesion |
| **CDH2** | N-cadherin | - | Cell-cell adhesion |

<sup>1</sup> Data from reference [49]; RGD and N3 are designations used in the database, and correspond to PIC-R and PIC-N, respectively.

Overall, qPCR analysis revealed distinct but partially overlapping transcriptional responses across the three hydrogel conditions (**Fig. 7**). Genes associated with ECM remodeling and mechanotransduction were consistently upregulated in PIC-R. In particular, *CYR61, FBLN2*, and *TSP1* showed significantly higher expression compared to both PIC-N and PIC-H, indicating enhanced matrix remodeling activity under integrin-mediated signaling under RGD functionalization (**Fig. 7**a-c). *MXRA5* followed a similar trend, although with a lower magnitude of change (**Fig. 7**d). These results support the activation of a mesenchymal-like transcriptional program in RGD-functionalized hydrogels.

**Figure 7.**
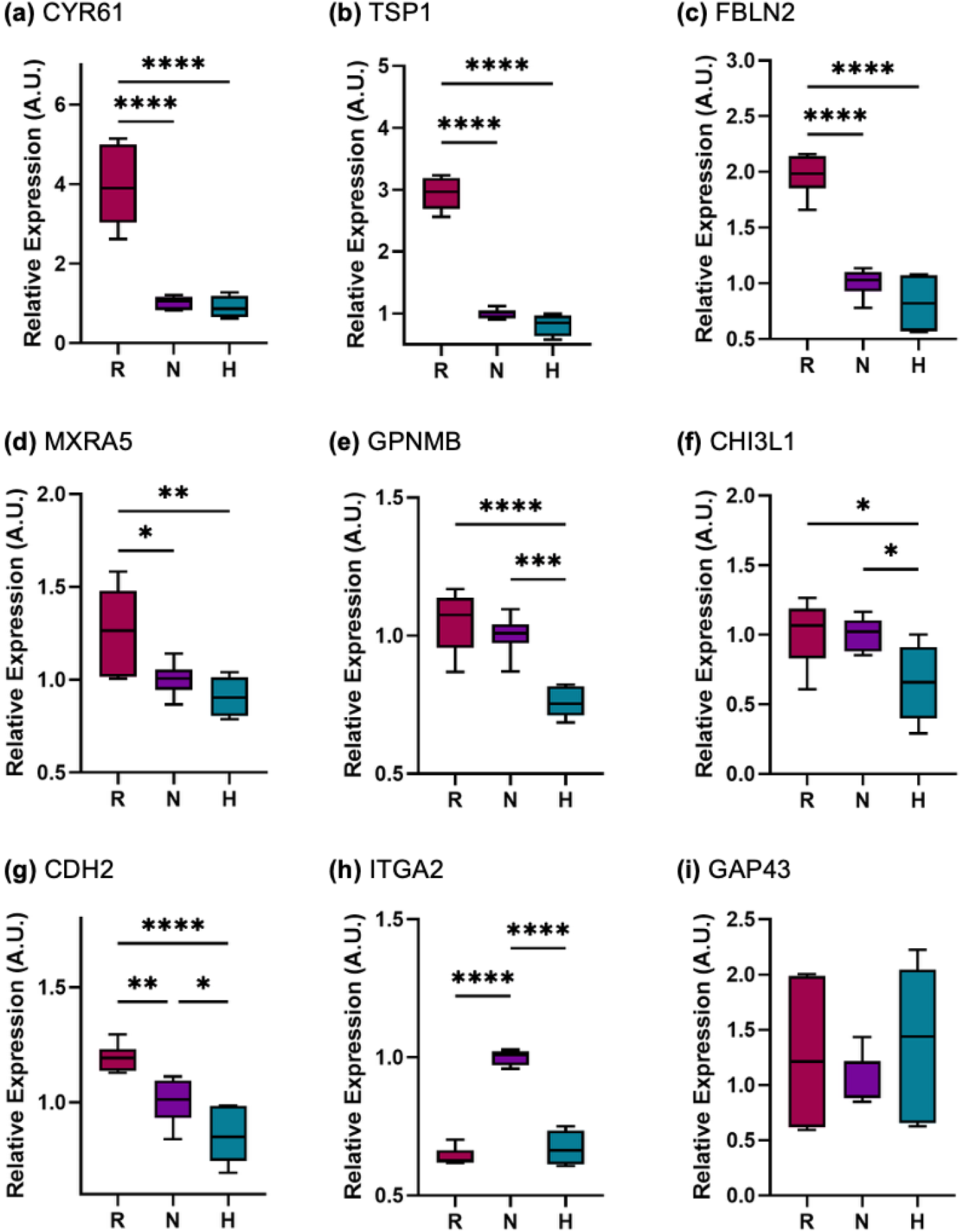
Quantitative PCR analysis of selected migration-related genes in cells cultured in three types of hydrogels for 48 h. Data are presented as mean ± SD of three independent biological replicates (each tested from technical triplicates). Result was normalized to group PIC-N. Differences between groups were analyzed using one-way ANOVA.

In contrast, PIC-N was characterized by increased expression of genes associated with cellular adaptation to low-adhesion environments. *ITGA2* was significantly upregulated in PIC-N compared to PIC-R and PIC-H, suggesting compensatory response to the limited integrin engagement (**Fig. 7**h). Other candidates identified in the proteomic dataset, including *CHI3L1* and *GAP43*, did not show significant enrichment at the mRNA level in the present system, which may reflect differences in matrix properties (e.g. PIC polymer length) and culture conditions relative to the reference proteomic study[49] (**Fig. 7**f,i).

Given the observed upregulation of *ITGA2* in PIC-N, which may reflect compensatory transcriptional responses under limited adhesion, we next examined whether cadherin expression was similarly regulated. qPCR analysis suggested that cadherin transcription may also be subject to compensatory regulation, as mRNA content was the lowest in PIC-H (**Fig. 7**g). In contrast, immunofluorescence analysis (**Fig. 3**m) revealed pronounced cadherin clustering in PIC-H hydrogels, suggesting that cadherin engagement in this condition is primarily regulated at the level of protein organization rather than transcription. More broadly, HAVDI-functionalized hydrogels did not display a distinct transcriptional profile comparable to either PIC-R or PIC-N. Instead, expression levels for most genes in PIC-H were similar to or intermediate between those observed in other conditions. Taken together with the enhanced cadherin clustering observed in PIC-H, these results suggest that the hybrid migration phenotype observed in this condition is not driven by a unique transcriptional program, but rather by the spatial organization and regulation of adhesion complexes.

Overall, these findings indicate that while mesenchymal and amoeboid-like migration modes are associated with distinct transcriptional signatures, the hybrid migration behavior observed in HAVDI-functionalized hydrogels is primarily at the level of adhesion architecture rather than large-scale transcriptional reprogramming. In combination with the observed differences in YAP nuclear localization, these results support a model in which ligand-dependent adhesion modulates how cells sense and respond to the mechanical properties of their surroundings, which in turn shape gene expression and ultimately define migration strategy.

## 4. Conclusion

In this study, we investigated how the biochemical cues present in synthetic 3D biomimetic environments influence single-cell migration modes and mechanotransduction in human adipose-derived stem cells. Using polyisocyanides (PIC) hydrogels functionalized with distinct adhesive ligands (RGD, HAVDI, or none), we identified three migration phenotypes associated with differential adhesion organization, force transmission, and YAP nuclear localization. RGD-functionalized matrices promoted integrin clustering, strong mechanical coupling to the matrix, extensive matrix remodeling, and a mesenchymal-like migration mode characterized by actin stress fiber formation and nuclear YAP enrichment. In contrast, non-adhesive matrices showed limited adhesion complex formation, minimal force transmission, and amoeboid-like migration with low YAP activation. HAVDI-functionalized matrices supported cadherin clustering and intermediate levels of mechanical coupling, leading to hybrid migration behaviors without a distinct transcriptional program.

Taken together, these findings demonstrate that ligand identity controls migration mode by regulating adhesion organization and downstream force transmission to the nucleus, even in matrices with comparable bulk mechanical properties. By linking adhesion cues to cytoskeletal organization, matrix remodeling, and mechanotransduction, this work establishes a direct mechanistic framework connecting extracellular ligand presentation to intracellular signaling and migration strategy. More broadly, our results highlight the potential of tunable synthetic matrices to dissect ligand-specific mechanobiological responses and provide a versatile platform to study migration plasticity in physiological and pathological contexts.

## Acknowledgements

The authors thank Dr. Egbert Oosterwijk for providing hASCs, and the developers of TFMLab for making the software publicly available and for its continued maintenance. We thank Dr. Charlotte Cresens for helping with immunofluorescence experiments. H.Z. acknowledges the financial support from the China Scholarship Council (Grant No. 202208440143). We acknowledge additional financial support from the National Natural Science Foundation of China (32471368), the Research Foundation of Flanders (FWO) research grant (G0C2422N) and postdoctoral fellowships (for G.S.: 12AML24N, for B.L.: 12AGZ24N, for H.Y: 12A2423N). This work also received financial support from KU Leuven (C14/22/085). Figure 1 was created with BioRender.

## CRediT authorship contribution statement

**Haoxiang Zhang:** Writing – original draft, Visualization, Methodology, Investigation, Formal analysis, Data curation, Validation, Software, Conceptualization. **Guillermo Solís Fernández:** Investigation, Formal analysis, Writing – review & editing. **Boris Louis:** Software, Writing – review & editing **Sarah Vorsselmans**: Writing – review & editing, Conceptualization. **Johan Hofkens, Paul Kouwer, Hongbo Yuan, and Susana Rocha**: Project administration, Supervision, Methodology, Resources, Funding acquisition, Conceptualization, Writing – review & editing.

