## Supporting Information for "Synthetic Fibrous Hydrogels as Minimal Systems to Modulate Cell Migration Modes in 3D"

**The supplementary information includes:**

**Table S1.** qPCR primer information

**Fig. S1.** Cell morphology in three PIC gels after 48 hours of culture.

**Fig. S2.** Cell Trajectories

**Fig. S3**, Quantitative analysis of Cadherin immunofluorescence.

**Fig. S4**, Quality control workflow for 3D YAP segmentation analysis.

**Fig. S5**, Analysis of cell migration turning angles.

**Fig. S6**, Persistence of cell migration over time.

**Table S1.** qPCR primer information

| Gene name | | sequence |
|---|---|---|
| *CYR61* | Fw | GGAAAAGGCAGCTCACTGAAGC |
| | Rv | GGAGATACCAGTTCCACAGGTC |
| *Thrombospondin-1 (TSP1)* | Fw | GCTGGAAATGTGGTGCTTGTCC |
| | Rv | CTCCATTGTGGTTGAAGCAGGC |
| *Fibulin-2* | Fw | CTGCTACAAGGCACTCACCTGT |
| | Rv | GTAGAAGGAGCCCTTGGTGTTC |
| *MXRA5* | Fw | TCACCGCTGAGACAGACACTGT |
| | Rv | CCGTTGTATCCTGGTATTCGGAG |
| *N-cadherin* | Fw | CCTCCAGAGTTTACTGCCATGAC |
| | Rv | GTAGGATCTCCGCCACTGATTC |
| *LRRC15* | Fw | CGTTGCTGTTCCAAGCGTCCAT |
| | Rv | GCTCAGTGGTAGAAGAGACGGA |
| *ITGA2* | Fw | TTGCGTGTGGACATCAGTCTGG |
| | Rv | GCTGGTATTTGTCGGACATCTAG |
| *GAP43* | Fw | GAGCAGCCAAGCTGAAGAGAAC |
| | Rv | GCCATTTCTTAGAGTTCAGGCATG |
| *Periplakin* | Fw | GGCTGCAGAATCTGGAGTTTGC |
| | Rv | CTCAGTCTCCTCATCCAGTTCC |
| *CHI3L1* | Fw | CCACAGTCCATAGAATCCTCGG |
| | Rv | TGCCTGTCCTTCAGGTACTGCA |
| *GPNMB* | Fw | GTGCTCAATGGAACCTTCAGCC |
| | Rv | AGGAATCCTACTCAGCTCCAGG |
| *KCTD12* | Fw | CTTCCGCTACATCCTGGATTACC |
| | Rv | AGCTCTGGCAGCTCGAAGTACT |
| *Perilipin-2* | Fw | GATGGCAGAGAACGGTGTGAAG |
| | Rv | CAGGCATAGGTATTGGCAACTGC |
| *RPL19* | Fw | CAGGCATATGGGCATAGGGAA |
| | Rev | TGCCTTCAGCTTGTGGATGT |

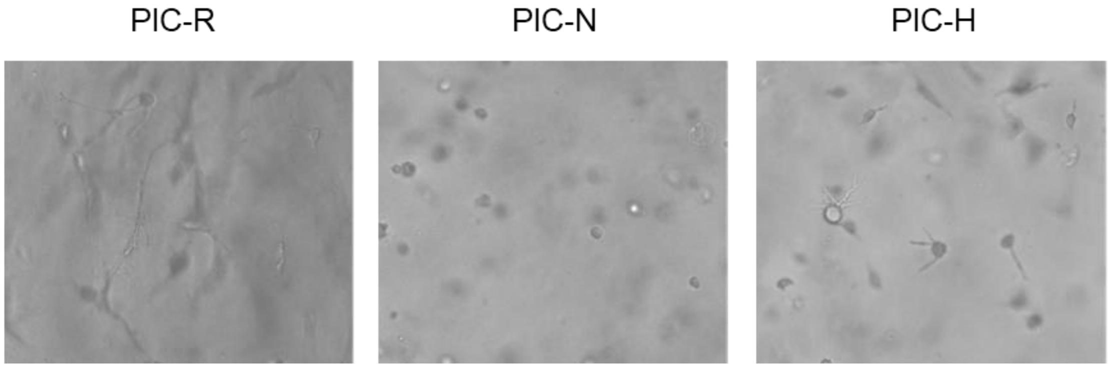


**Figure S1**, Cell morphology in three PIC gels after 48 hours of culture. Cells cultured in PIC-R gels exhibit an elongated, spreading morphology, while cells in PIC-N gels maintain a spherical shape. Cells in PIC-H gels show limited spreading capability. Scalebars = 50μm

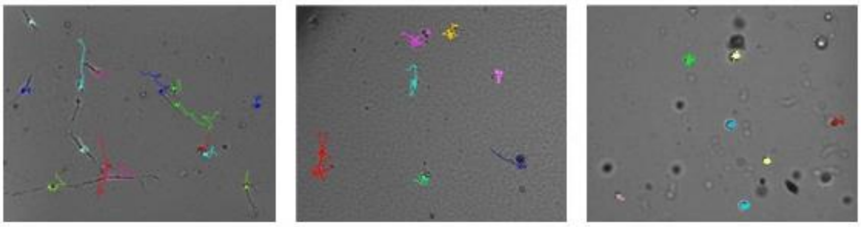

**Figure S2**, Trajectories were generated using the Manual Tracking plugin in ImageJ/Fiji and are overlaid on the final time point of the corresponding live-cell imaging sequence.

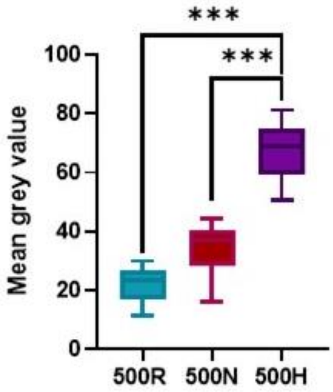


**Figure S3**, Quantitative analysis of Cadherin immunofluorescence. Mean gray values were measured within segmented cell areas to assess expression levels.

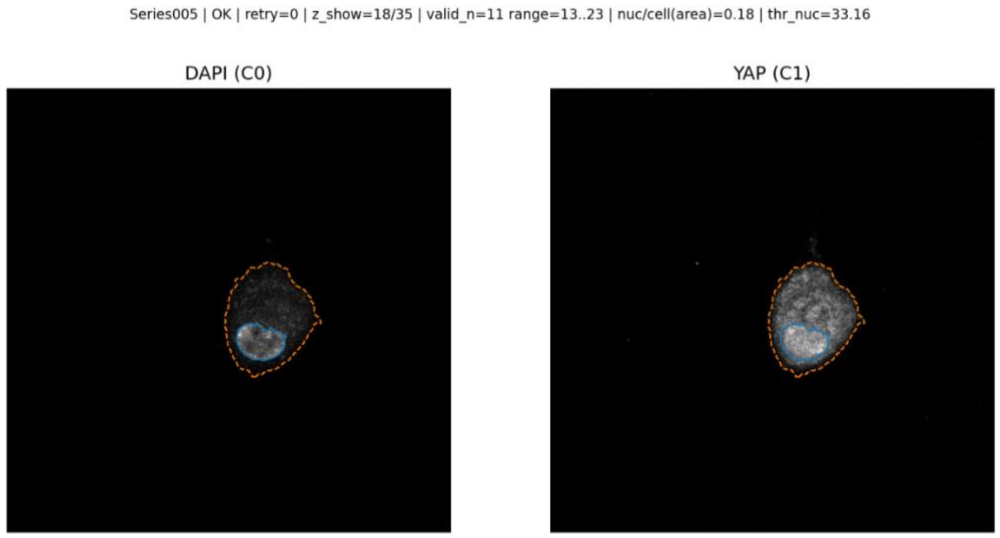


**Figure S4**, Quality control workflow for 3D YAP segmentation analysis. The process allows for parameter verification identical to batch analysis, generates visual overlays of segmentation results for manual validation against cellular and nuclear boundaries, and outputs a corresponding CSV file containing detailed segmentation metrics.

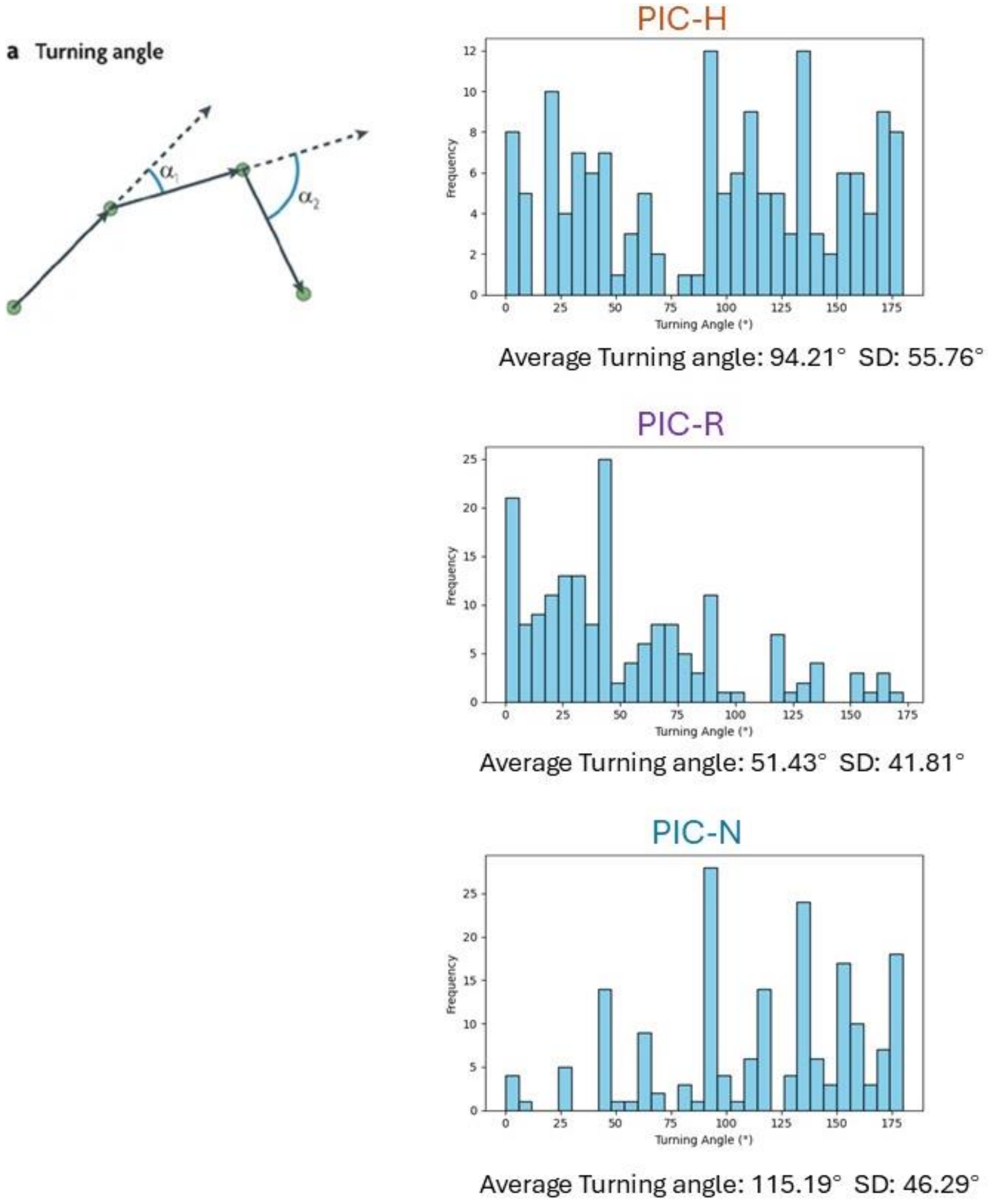


**Figure S5**, Analysis of cell migration turning angles. **a**, Schematic defining the turning angle between movement steps; **b**, Histogram showing the distribution of measured turning angles.

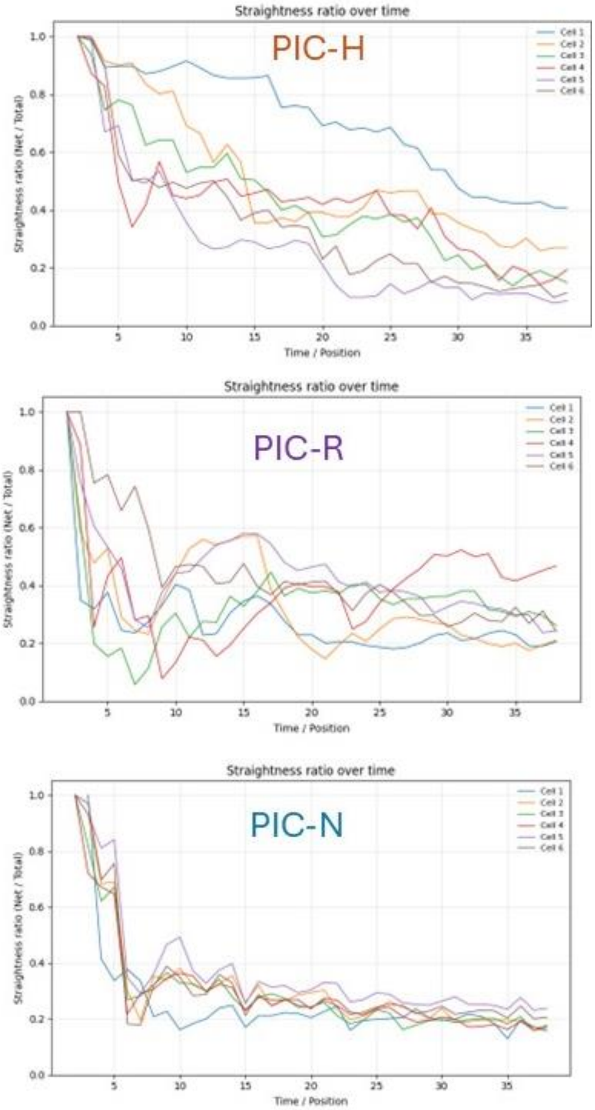


**Figure S6**, Persistence of cell migration over time. The ratio of net-to-total distance was calculated for each time point. Net distance is defined as the displacement from the starting position, while total distance represents the cumulative path length traveled.

SI movie/gif. (Will be separately upload when submit to the journal)

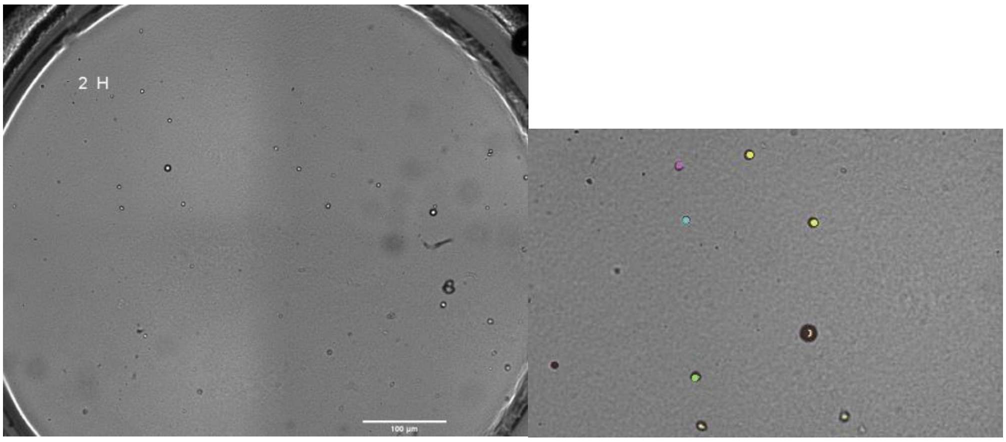


PIC-H Time-lapse images were acquired every 2 hours for over 72 hours.

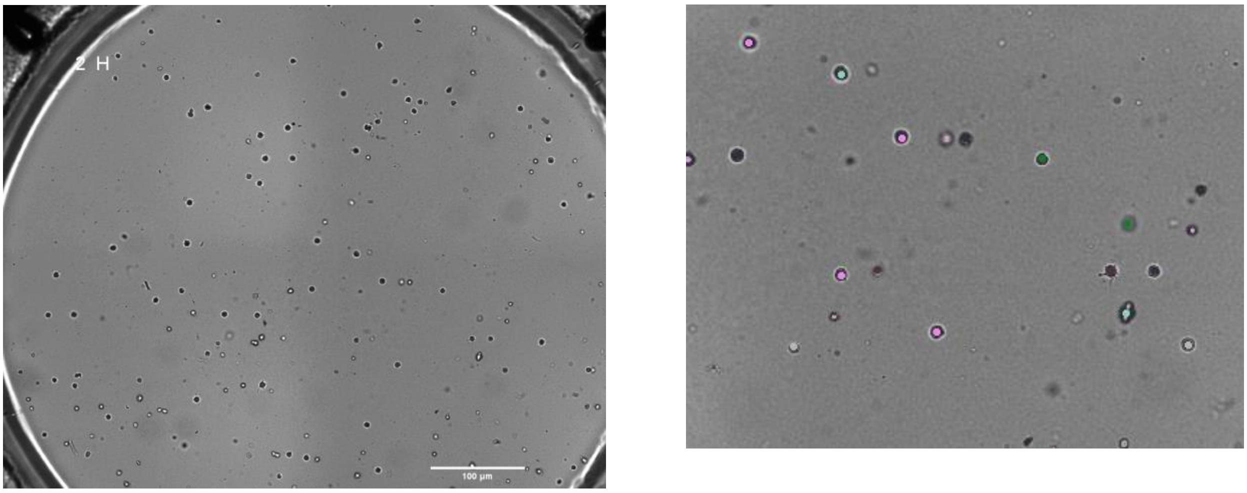


PIC-R

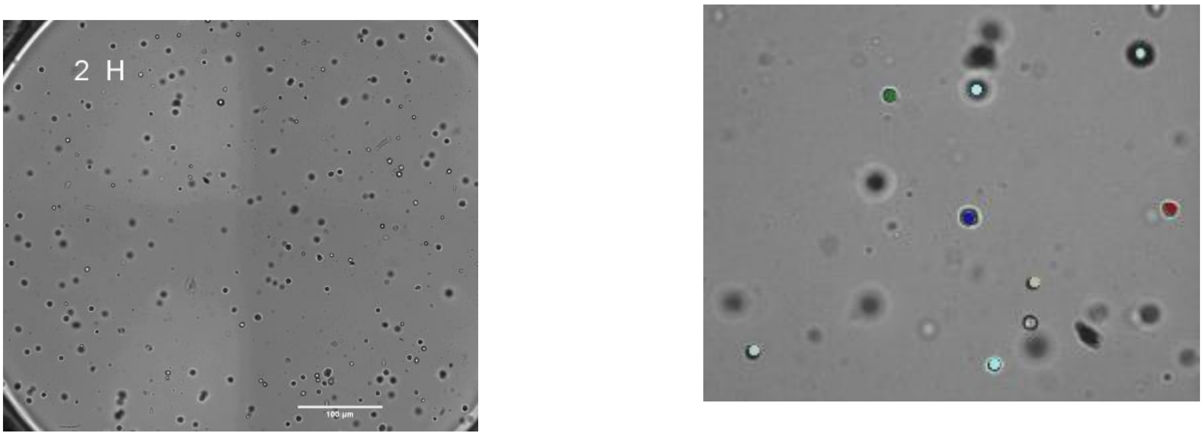


PIC-N
